# Dormant *Mycobacterium tuberculosis* reactivates *ex vivo* in blood from HIV infected individuals

**DOI:** 10.1101/2023.11.18.566455

**Authors:** Madhur Kalyan, Stuti Agarwal, Aman Sharma, Indu Verma

**Affiliations:** Department of Biochemistry, Post Graduate Institute of Medical Education and Research, Chandigarh, India; Department of Biochemistry, All India Institute of Medical Sciences, Jammu, India; Division of Pulmonary, Allergy and Critical Care Medicine, Stanford University, Stanford, California, USA; Department of Internal Medicine, Post Graduate Institute of Medical Education and Research, Chandigarh-160012, India

**Keywords:** Tuberculosis, HIV infection, reactivation, CD4 counts, dormant *Mycobacterium tuberculosis*

## Abstract

Reports of tuberculosis in Human Immunodeficiency Virus infected patients despite high CD4^+^ T-cell counts suggest different mechanisms of reactivation. *In vitro* developed dormant *Mycobacterium tuberculosis* H37Rv were cultured in HIV blood and normal blood. Colony forming units analysis was performed and expression of mycobacterial Resuscitation Promomoting Factor genes was analyzed by qRT-PCR. The *in vitro* developed dormant *M. tb* resuscitated in HIV blood as evident from increase in CFUs and up-regulation in the expression of mycobacterial *rpf* genes. Reactivation of dormant tubercle bacilli following HIV infection in spite of high CD4^+^ T-cell counts could be due to some viral or host factors which needs further studies.

## 1. Introduction

Tuberculosis, caused by *Mycobacterium tuberculosis* (*M. tb*), has become a leading killer alongside HIV. According to World Health Organisation statistics, approximately 10 million people fell ill with tuberculosis (TB) worldwide and 1.4 million people lost their lives to TB in 2019 (together with 208000 people with HIV) [1]. Inpeople living with HIV, tuberculosis is one of the leading causes of death along with greater risk of progression from latent to active TB as compared to those without HIV infection.There is a profound effect of HIV infection on immune system functions anda significant contributor to immune suppression is the depletion of CD4^+^T cells [2]. The result of this immune suppression is just like a blessing for opportunistic infections (OIs) [3] and among these OIs, tuberculosis is one of the major causes of mortality among HIV infected individuals. Risk of acquiring tuberculosis infection increases exponentially with decline in the CD4^+^T-cells as the HIV infection progresses [4]. Although immune suppression, in particular, due to a decline in CD4 count, seems to be the major cause of occurrence of tuberculosis, there are reports that suggest TB infection is contacted even when the CD4 count is intact [5]. It is known that in HIV infection, irrespective of CD4^+^ T cell counts and viral load, there is much higher risk for TB [6]. Recently a study on the non-human primates, infected with Simian Immunodeficiency Virus, also suggested that only CD4 ^+^ T cells depletion during latent TB infection was not sufficient to reactivate latent TB suggesting different mechanism of TB reactivation at higher CD4^+^ T cell numbers in HIV infected individuals [7]. These alternative mechanisms may be dependent on various mycobacterial, HIV and host factors and interplay of these factors may be responsible for the reactivation of latent TB in HIV infected individuals. It is well documented that, as a part of its survival strategy, *M. tb* is able to adapt to various host physiological environments and it can transit between latent and active states inside the host cells. Recently, it has been demonstrated that *M. tb* can differentiate between diverse blood environments of HIV patients and healthy individuals [6]. In addition, it has been proposed that *M. tb*, for its optimal growth, alters its gene expression in the altered immunological environment in the blood of HIV patients [6]. A variety of genes have been reported to be expressed in mycobacteria, which are responsible for maintaining its dormancy, but not much was known about the genes responsible for growth and resuscitation of dormant mycobacteria until the discovery of resuscitation promoting factors (Rpfs) in *Micrococcus luteus* [8][9]. These factors are able to resuscitate the bacteria from a non-culturable state at picomolar concentrations [9]. There are five Rpf homologues (Rpf A-E) in *M. tb* that share sequence as well as functional homology with Rpf proteins of *M. luteus* and are expressed during different phases of growth and resuscitation [10][11]. In *M. tb*, RpfB, a lytic transglycosylase has been reported to interact with an Rpf interacting protein A (RipA), the D,L-endopeptidase that empowers these proteins to harmonically degrade peptidoglycan, a critical process for cell growth and division [12].

The transition of latent TB to active TB in HIV infected individuals, specially at higher CD4 counts, could therefore be due to the mycobacterial resuscitation mediated either by the virus itself or through some unknown host factors which ultimately may lead to change in gene expression of mycobacterial resuscitation promoting factors for the process of “reawakening” of dormant bacilli. Hence, we studied the reactivation of dormant *M. tb* in blood microenvironment from HIV patients with CD4 counts ≥350 cells/μl.

## 2. Material and Methods

### 2.1. Bacterial cultures

*Mycobacterium tuberculosis* H37Rv (NCTC 7416) was cultured and maintained on Sauton’smedium and bacterial aliquots were stored at -80ºC for further usein various experiments.

### 2.2. Human subjects

Peripheral blood samples from six HIV infected and six healthy individuals were collected with written informed consent. HIV positive individuals (CD4≥350 cells/μl) for the study were recruited from the Department of Internal Medicine, PGIMER, Chandigarh, India. It included Antiretroviral therapy (ART)-naive asymptomatic HIV patients tested positive for HIV infection based on rapid HIV screening test i.e., Coombs AIDS, Tridot and confirmed by HIV ELISA (as per guidelines of the National AIDS Control Organization, India) without any clinical, radiological and bacteriological evidence of pulmonary/extrapulmonary tuberculosis at the time of recruitment. Ethical clearance for the study was granted by Institute Ethical committee, PGIMER, Chandigarh, India (8823-PG 11-1 TRG/12484) and samples were collected after taking the written informed consent.

### 2.3. Development of dormant M. tb models

#### 2.3.1. Long term hypoxia induced model of dormancy

The method used to develop dormant *M. tb* model has been adopted from the study of Shleeva et al. [13], in which *M. tb* has been cultured under anaerobic conditions for an extended period of 8 months, that correlates to hypoxic conditions, and hence it is referred as long term hypoxia induced dormant *M. tb* model in this study. For development of this model, 1x10^6^ cells/ml of log phase *M. tb* were inoculated into 135ml Sauton’s media supplemented with ADC (15ml) and 0.05% tween-80 in a sealed 500 ml flask and cultured at 37ºC with shaking at 170 rpm for 7 months [13]. At selected intervals of 30 days, cell viability was evaluated by carrying out colony forming unit (CFU) assay. The hypoxia induced dormant *M. tb* obtained at the end of 7 months wasused for further experiments.

#### 2.3.2. Development of potassium deficiency induced dormant M. tb model

Potassium deficient model of *M. tb* H37Rv was developed as previously described [14]. Briefly, to induce dormancy and obtain dormant bacilli, *M. tb* culture was grown for 22-25 days with shaking at 170 rpm at 37ºC to obtain a culture rich in stationary phase bacilli. Stationary phase *M. tb* thus obtained wasinoculated into 100 ml potassium deficient Sauton’s media supplemented with ADC and containing 0.05% tween-80 at 5x10^5^ cells/ml and grown in shaking conditions at 170 rpm at 37ºC for 30 days. Thirty-day starved culture was further incubated for another 15 days in the presence of rifampicin at a concentration of 5μg/ml to obtain a maximal dormant cell population.

### 2.4. Resuscitation of dormant M. tb

#### 2.4.1. Culturing dormant M. tb bacilli in HIV and Normal blood

Dormant *M. tb* were cultured in blood (HIV or normal) as previously described [15] with modifications. Briefly, venous blood samples collected from HIV infected and normal individuals werediluted with equal volume of RPMI-1640 medium containing 10mM HEPES and 1mM glutamine. Single cell suspensions of dormant *M. tb* (0.1ml) were inoculated with 1:1 diluted blood (0.9 ml) in 7ml endotoxin free Sterilin Bijou tubes (Dynalab corporation, USA) and cultured for 4 days at 37ºC with slow shaking at 80 rpm.

### 2.5. Colony forming unit enumeration of dormant M. tb grown in HIV and normal blood

For CFU assay, blood cells from 4 day old cultures were lysed with 0.1% Triton X-100 in water for 15 min at room temperature followed by centrifugation at 2500xg for 15 min. Pellets thus obtained were washed with fresh Sauton’s media and finally dispensed in 1 ml Sauton’s media. Serial dilutions were prepared and CFU assay was performed as described earlier [6] at day zero and 4 after culture in HIV/Normal blood.

### 2.6. RNA isolation and quantitative real time RT-PCR

Briefly,bacterial suspensions from both the dormancy models as well as from dormant *M. tb* cultured in HIV and normal blood were transferred to o-ring bead beater tubes and centrifuged for 10 minutes at 2500 rpm, pellet was resuspended in Trizol solution (ThermoFisher Scientific, USA.) and RNA isolation was carried out as per manufacturer’s instructions. RNA samples were DNase treated as per manufacture’s protocol using DNase I, RNase free kit (ThermoFisher Scientific, USA.) and Verso cDNA synthesis kit (Thermo Fisher Scientific, USA.) was used for first strand cDNA synthesis. Fold change in the expression of specified genes (*rpf/ripA/hspX*) was analyzed for dormant *M. tb* (hypoxia and potassium deficiency) with respect to log phase active bacilli whereas for dormant bacilli cultured in HIV blood with respect to those cultured in normal blood. Quantitative real time RT-PCR was performed on the Qiagen Rotor Gene Q Real Time PCR machine by using SYBR chemistry. 16s rRNA was used as housekeeping gene for normalizing the transcript levels of respective genes. Melt curve analysis was performed for each gene to ensure that fluorescence levels detected were due to the amplification of a specific gene. Sequences of the primers for respective genes are as follows: *rpf A* (Forward): CCACTGGCTTCGGGTGTTA, *rpf A* (Reverse): CCAGCGGTGCGGGCAGGTCGTTAG; *rpf B* (Forward): CGACGCTAAGCAGGTGTGGACGAC, *rpf B* (Reverse) : CACTCAGCAGCCCCGCGACATTGG; *rpf C* (Forward): GTCACGGCATCCATGTCGCTCTCC, *rpf C* (Reverse): CCCAGGTGGCCGGCTTGAACT; *rpfD* (Forward): GCCGCGAGTCCCCAGCAACAGAT, *rpfD*(Reverse): GGCCGCGAGGAACGTCAGGATG; *rpfE*(Forward): CCAGCCGGTATCGCCAATG, *rpfE*(Reverse): CCACCGGACTCGCACTG; *ripA*(Forward): CACCGATACCGGCATCACCA, *ripA*(Reverse): AGGGCACCCCGATCTGTGAC; *hspX*(Forward): CAAATGGCCACCACCCTTCC, *hspX*(Reverse): CTCGTCTTCCAGCCGCATCA; *16s* (Forward): TCCCGGGCCTTGTACACA, *16s* (Reverse): CCACTGGCTTCGGGTGTTA

Approximately 25ng RNA was used for qRT-PCR analysis. Real-time PCR analysis was performed using the following conditions: 95ºC for 10 min–initial denaturation, 40 cycles of denaturation at 95ºC for 10 s, annealing at 60ºC for 20s and extension at 72ºC for 30s. Melt curve was also run at the end of amplification curve. The C_t_ values obtained for every sample was used to calculate relative fold change by using delta-delta C_t_ method and change in expression was represented as log_2_ fold change with respect to controls.

## 3. Results

### 3.1. Hypoxic and potassium deficient environment restricts growth of M. tb in vitro

Hypoxia induced *in vitro* model of *M. tb* dormancy was developed by culturing in sealed flasks containing Sauton’s media for a period of seven months and cell viability was analysed by cfu assay at selected intervals of 30 days. Fig. 1a shows the log10 cfu counts from time point zero to month seven. Once it was established that cfu counts do not change after fourth month onwards, experiment was repeated thrice to confirm that there is no change in cfu counts after fifth and seventh month. Fig. 1b shows log_10_ cfu/ml values at fifth (4.64±0.02) and seventh month (4.63±0.01) with no significant change in cfu numbers thus confirming the presence of non-dividing bacilli in the culture. In case of bacilli cultured under potassium deficient conditions, the observed decrease in log_10_ cfu/ml for dormant *M. tb* cultured in potassium limiting conditions was significant as compared to logarithmically grown *M. tb* H37Rv (Fig. 1c).

**Fig. 1.**
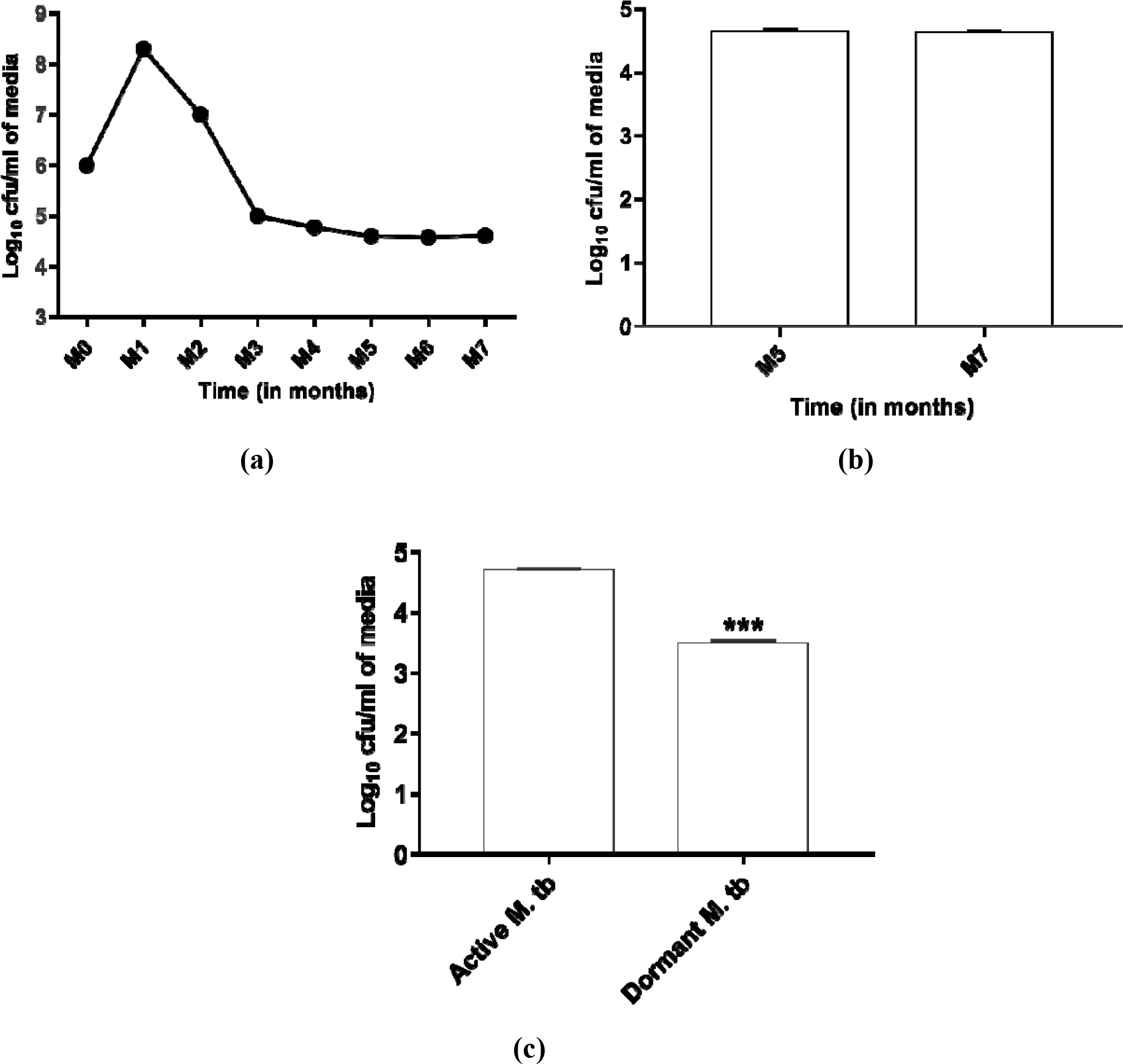
Development of models of dormant *Mycobactrium tuberculosis* H37Rv. (a) Changes in log10 cfu/ml every 30 days from month zero up to month 7 (M0 to M7) in *M. tb* cultured under hypoxic conditions. Each dot on the graph represents the means of three technical replicates at each time point. (b) Confirmation of establishment of dormancy based on changes in log10 cfu/ml at month 5 (M5) and month 7 (M7). (c) Changes in log10 cfu/ml of potassium deficiency induced dormant *M. tb* with respect to logarithmically grown active *M. tb*. Each bar represents the mean ± SD of log10 cfu/ml from three independent experiments. p (***) <0.001 w.r.t. logarithmically grown *M. tb* H37Rv by independent t-test.

Further confirmation of dormancy was also established at the gene level in both the models. In case of hypoxia model, relative mRNA expression of dormant *M. tb* was studied at fifth and seventh months of dormancy with respect to actively growing log phase bacilli in Sauton’s growth medium (Fig. 2a,2b). The dormancy associated gene *hspX* achieved a significant up-regulation at fifth month (p<0.05) with a log_2_ fold change of 1.27±0.08 (Fig. 2a) which increased further at month 7 (p<0.05) with a 2.5±0.53 log_2_ fold change with respect to log phase actively growing bacilli(Fig. 2b). A significant decrease in the relative expression of *rpfA*(p<0.01), *rpfB* (p<0.01), *rpfD* (p<0.001), *rpfE* (p<0.05) and *ripA* (p<0.05) gene was observed at fifth and seventh month(Fig. 2a,2b). Similar to the hypoxia induced model, changes in the *rpf* genes expression were observed for the bacilli cultured under potassium deficient conditions with a significant reduction in the mRNA expression of all the *rpf* genes (Fig. 2c).

**Fig. 2.**
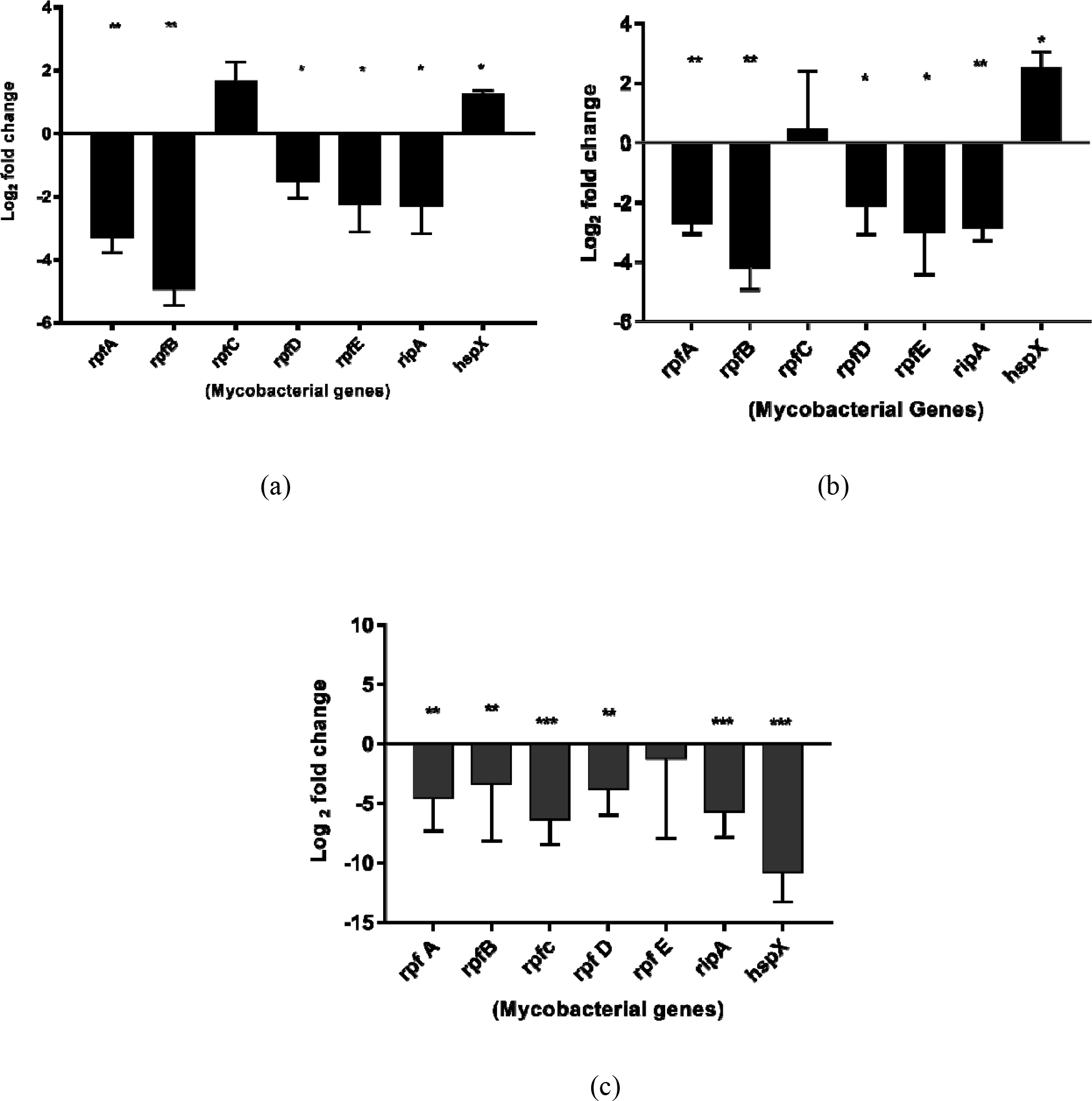
Changes in the expression of five *rpf*genes (*A-E*), *ripA*and *hspX*genes during the establishment of dormancy. Log2 fold change in the expression of various genes in **(a)** hypoxia model at month 5 **(b)** hypoxia model at month 7 and **(c)** potassium deficiency induced dormant bacilli. Upregulation is indicated by log2 fold change (Y-axis values) of ≥1 and <-1 indicates down-regulation. Each bar depicts mean ± SD values for each of the genes from three independent experiments. p(*)<0.05, p(**)<0.01 w.r.t log phase *M. tb* H37Rv (independent t-test). For relative expression analysis, RNA isolated from log phase *M. tb* H37Rv culture was used as positive calibrator and for normalization 16s rRNA was used as reference gene.

### 3.2. Hypoxia induced dormant M. tb reactivates ex vivo in HIV blood

At the end of the 4 days of incubation period, in comparison to the dormant *M. tb* (hypoxia induced) at day zero (4.66±0.01 log_10_ cfu/ml), there was a significant increase in the cfu counts of *M. tb* cultured in the HIV blood (5.55±0.006 log_10_cfu/ml) (p<0.001) whereas a significant decrease of those cultured in normal blood (4.31±0.04 log_10_cfu/ml) (p<0.001) was observed (Fig. 3a). Fig. 3a also shows a significant increase in the log_10_cfu/ml for the bacilli cultured in HIV blood (p<0.001) as compared to those cultured in normal blood. This increase in cfu following culture in HIV blood even with high CD4 counts indicated the reactivation of *M. tb* in such an environment. Further, at the end of the incubation period, with respect to hypoxic dormant *M. tb* cultured in normal blood, a significant up-regulation in the expression of *rpfA, rpfB, rpfC, rpfE and ripA* genes and a significant downregulation for *hspX* (p<0.001) gene was observed for those cultured in HIV blood (Fig. 3b). Additionally, the mRNA expression of *hspX* gene following reactivation in HIV blood was decreased with a 1.36±0.29 log_2_ fold reduction with respect to hypoxic dormant *M. tb* cultured in normal blood (Fig. 3b).

**Fig. 3.**
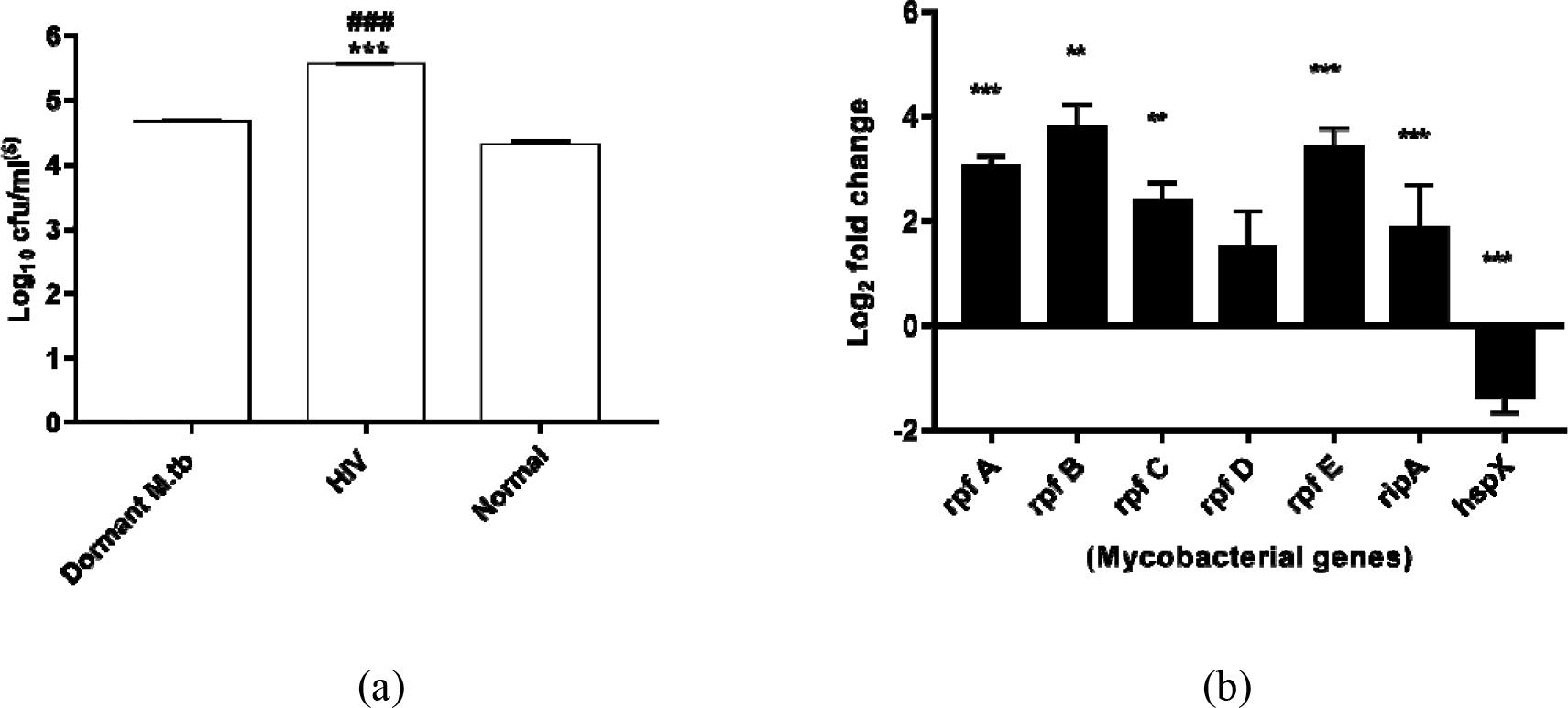
Effect of HIV and normal blood on the resuscitation of hypoxia induced dormant *M. tb* H37Rv. Hypoxia induced dormant *M. tb* was cultured in HIV and normal blood for cfu enumeration and mRNA expression of various genes at the end of 4 days incubation. **(a)** Changes in cfu of hypoxia induced dormant *M. tb* cultured in the presence of either HIV or normal blood. Each bar represents the mean±SD of log10 cfu/ml from three independent experiments. p(***) <0.001 w.r.t hypoxia induced dormant *M. tb* at day zero, p(###) <0.001 w.r.t hypoxia induced dormant *M. tb* cultured in normal blood at day 4 (Post hoc Bonferroni) (**$)** indicates log10 cfu/ml of media in case of dormant *M. tb* at day zero and log10 cfu/ml of blood in case of HIV/normal blood at day 4. **(b)** Log2 fold changes in the expression of five *rpf*genes (*A-E*), *ripA*and *hspX*genes in dormant *M. tb* cultured in HIV blood as compared to those cultured in normal blood. For relative expression analysis, RNA isolated from hypoxia induced dormant *M. tb* cultured in normal blood culture was used as positive calibrator for and for normalization 16s rRNA was used as reference gene. Upregulation is indicated by log2 fold change (Y-axis values) of ≥1 and <-1 indicates down-regulation. Each bar represents mean ± SD values for each of the genes from three independent experiments. p(**) <0.01, p(***) <0.001w.r.t hypoxia induced dormant *M. tb* cultured in normal blood (independent t-test).

### 3.3. Reactivation of potassium deficient dormant bacilli in HIV blood

Potassium deficient dormant *M. tb* resumed growth following culture in HIV blood at the end of the incubation period of four days as evident from a significant (p<0.001) rise in cfuof 8.4±0.45 x 10^3^ cfu/ml from 3.13± 0.27 x 10^3^cfu/ml of potassium deficient dormant *M. tb* at day zero (Fig. 4a). A similar phenomenon of resuscitation was not observed for dormant *M. tb* cultured in normal blood since no significant change in cfu was observed in comparison with the potassium deficient dormant *M. tb* at day zero. On comparison of growth of potassium deficient dormant *M. tb* between those cultured in HIV blood with respect to normal blood, it was observed that colony forming units for the former (8.4±0.45 x 10^3^cfu/ml) were significantly increased (p<0.001) with respect to the latter (3.76±1.3x 10^3^cfu/ml) (Fig.4a). Resuscitated bacilli from HIV blood showed a significant up-regulation in relative mRNA expression of *rpfA*(p<0.05) and *rpfB*(p<0.05)genes, with respective log_2_ fold changes of 5.2±0.8 and 7.6±0.3 when compared to potassium deficient dormant *M. tb* cultured in normal blood. Such a significant fold change was not observed for *rpfC, rpfE, ripA* and *hspX*genes rather a significant reduction (p<0.001) in the *rpfD* gene was observed (Fig.4b)

**Fig. 4.**
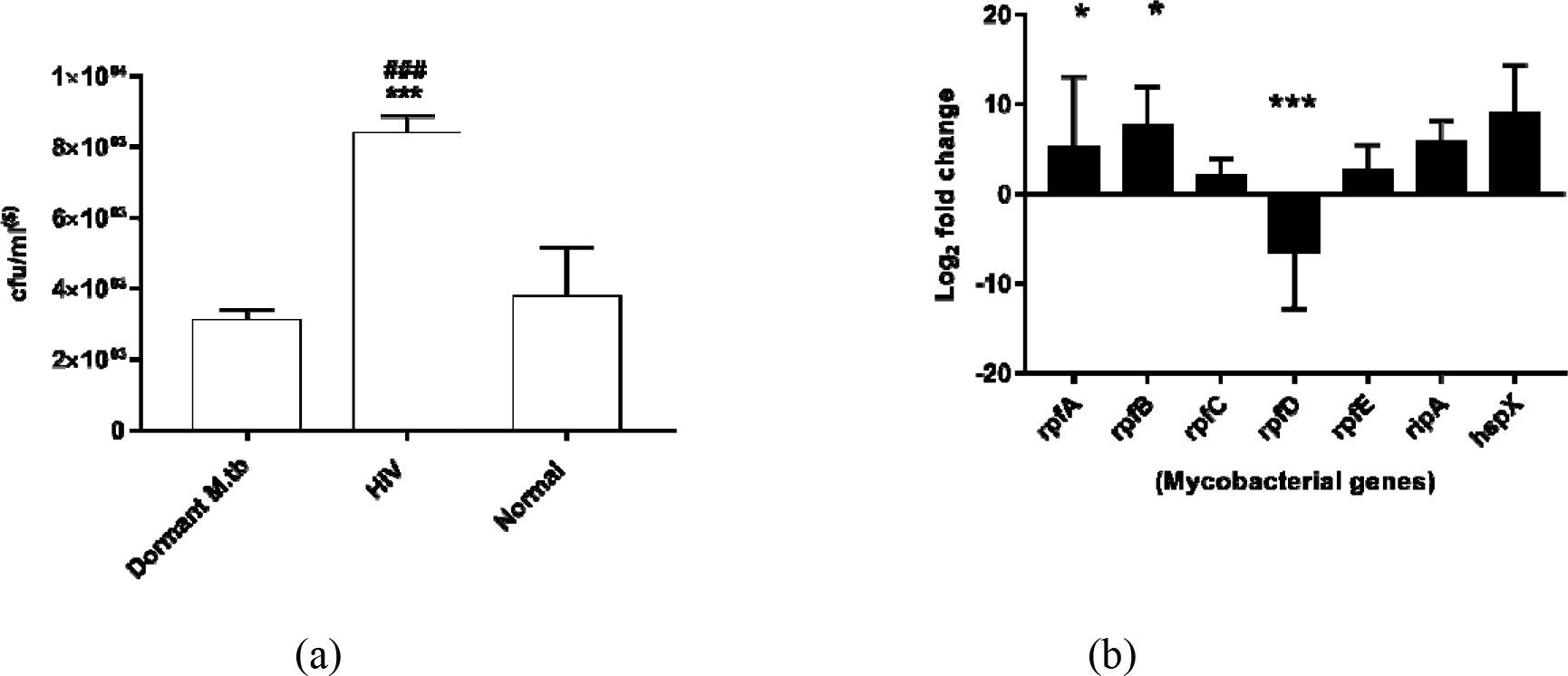
Effect of HIV and normal blood on the resuscitation of potassium deficiency induced dormant *M. tb* H37Rv. Potassium deficiency induced dormant *M. tb* was cultured in HIV and normal blood for cfu enumeration and mRNA expression of various genes at the end of 4 days incubation. **(a)** Changes in cfu of potassium deficiency induced dormant *M. tb* cultured in the presence of either HIV or normal blood. Each bar represents the mean±SD of log10 cfu/ml from three independent experiments. p(***) <0.001 w.r.t Potassium deficiency induced dormant *M. tb* at day zero, p(###) <0.001 w.r.t Potassium deficiency induced dormant *M. tb* cultured in normal blood at day 4 (Post hoc Bonferroni) (**$)** indicates log10 cfu/ml of media in case of dormant *M. tb* at day zero and log10 cfu/ml of blood in case of HIV/normal blood at day 4. **(b)** Log2 fold changes in the expression of five *rpf*genes (*A-E*), *ripA*and *hspX*genes in potassium deficiency induced dormant *M. tb* cultured in HIV blood as compared to those cultured in normal blood. For relative expression, analysis RNA isolated from potassium deficiency induced dormant *M. tb* cultured in normal blood was used as positive calibrator and for normalization 16s rRNA was used as reference gene. Upregulation is indicated by log2 fold change (Y-axis values) of ≥1 and <-1 indicates down-regulation. Each bar represents mean ± SD values for each of the genes from three independent experiments. p(*) <0.05, p(***) <0.001w.r.t potassium deficiency induced dormant *M. tb* cultured in normal blood (independent t-test).

These results clearly suggest that dormant mycobacterial bacilli are able to reactivate in the microenvirnment of HIV blood.

## 4. Discussion

The success of *Mycobacterium tuberculosis (M. tb)*, causative agent of tuberculosis, is primarily attributed to its potential to enter into a state of dormancy. Competence of immune system reduces in HIV infection and it has also been seen as a potent predisposing factor for the reactivation of tuberculosis [16]. Reactivation of tuberculosis following HIV infection has been attributed to decline in CD4^+^ T-cells[17][18][19][20], however, data from previous human and animal studies have demonstrated that decline in CD4^+^ T-cells does not correlate directly to occurrence of TB [7].This study was therefore planned to study the reactivation of dormant tubercle bacilli by culturing in blood microenvironment from HIV patients with CD4 counts ≥350 cells/μl. Considering the scarce information available on the *in vivo* conditions prevailing inside host during latent infection and no single model of dormancy can mimic all the conditions that might prevail, two different *in vitro* models of dormancy were used in the current study. In long term hypoxia model, a consistent cfu was obtained post four months up to seven months with no significant change in the cfu numbers (Fig. 1), suggesting that bacilli were in a viable non-dividing state, a hallmark of *M. tb* dormancy. Sufficient evidence is available in the literature for the persistence of *M. tb* during dormancy under *in vitro* nutrient starvation and oxygen deprivation conditions[21][22]. In our study, absolute non-culturability, defined by zero cfu, could not be attained as described by Shleeva et al.[13]. However, similar results were obtained in an earlier study where absolute non-culturability was not achievable in 80 to 100 days old stationary cultures [23]. Further, at molecular level also, in long term hypoxia model downregulation of reactivation associated genes along with upregulation of dormancy associated *hspX* gene [24][25] indicate towards achievement of non-replicating persistence. On the other hand, in case of potassium deficient dormancy model, there was significant reduction in cfu counts (Fig. 1) as supported by earlier studies [14] as well as decreased expression of *rpfA, rpfB, rpfC and rpfD* and *hspX* genes with respect to the log phase bacilli (Fig. 2). Downregulation in the expression of *hspX* gene in potassium deficient dormancy model could be explained on the basis that it is expressed in response to hypoxic conditions whereas in this model aerated cultures were used. The dormant *M. tb* obtained from both the models were further used to study the effect of HIV on resuscitation of these bacilli.

In order to study the effect of HIV on the resuscitation, dormant *M. tb* were cultured in blood from HIV infected donors with CD4 counts more than 350 cells/μl. Earlier, it has been well established that active *M. tb* can replicate in the blood environment[6][15][26].The replication of BCG has also been shown to be increased in whole blood from HIV positive children [27]. The present study is the first of its kind where blood environment was selected to study the resuscitation of dormant *M. tb*. Resuscitation could be observed when dormant *M. tb*, from both the models, were cultured in HIV blood as observed by increase in the cfu counts with respect to those cultured in normal blood. Active *M. tb* has been shown to adapt and survive in both HIV+ and HIV-blood [6], suggesting that dormant *M. tb* may also have a different transcriptional profile as compared to active bacilli in blood environment. At transcriptional level, in both the models, the relative expression of resuscitation genes involved in reactivation of mycobacteriawas up-regulated when cultured in HIV blood with respect to those cultured in normal blood (Fig. 3,4). The expression of dormancy associated gene *hspX* was stifled following resuscitation of hypoxia induced dormant bacilli in HIV blood with respect to those cultured in normal blood. Suppressed expression of dormancy related transcriptome and hence *hspX* was also observed in a recent study following culture of actively growing *M. tb*in HIV blood [6]. In the same study, increased expression of *rpfB* gene has been observed in the actively growing *M. tb* cultured in HIV blood further supporting the observations of present study that resuscitated bacilli resumed active replication in HIV blood. Resuscitation of dormant *M. tb* in HIV blood inspite of high CD4 counts raised the possibility of involvement of some other factors being involved in the process of resuscitation.

Some studies have correlated high HIV load in the tissues to be a cause of functional disturbance in granuloma [28][29][30]. HIV gp120 and p24 have been reported to be high during earlier stages of HIV infection [31][32] and also risk of TB reactivation is elevated during early HIV infection [20]. It is possible that some of viral proteins are contributing to reactivation of TB during HIV infection. A previous study showed that Tat protein from HIV can bind directly to *M. avium* through its integrin α5β1 present on its surface[33]. Presence of such bacterial integrins on the surface of *M. tuberculosis* that could interact with HIV proteins is not documented as yet.

In our study we have shown that inspite of a higher CD4 cell count dormant bacilli could resuscitate in blood from HIV infected donors. This scenario suggests some other intricate mechanisms responsible for the phenomenon involving viral or host factors. For instance, various host factors including cytokines are known to play a significant role in the progression and pathogenesis of HIV infection [34]. Interestingly, cytokines such as IL-1β, IL-6, and TNF-α have also been demonstrated to promote the growth of mycobacteria as well as other bacteria under *in vitro* conditions [35][36][37][38]. However, it is yet to be determined if the host cytokines can affect the reactivation of latent tubercle bacilli. The transition of latent TB to active TB in HIV infected individuals could therefore be due to the mycobacterial resuscitation mediated either by the virus itself or through host factors which ultimately may lead to change in gene expression of mycobacterial resuscitation promoting factors for the process of “reawakening” of dormant bacilli.

This study has some limitations, firstly, we could not achieve absolute non-culturability in our dormancy models. This infers that there still might be a small population of active bacilli along with the dormant bacilli in the culture. Hence, the colonies that appeared on the solid media, following resuscitation, might be a mixture of resuscited as well as active bacilli.

In conclusion, this study demonstrates that dormant *Mycobacterium tuberculosi s*from two different in *vitro* models could be resuscitated when cultured in HIV+ blood as evident from increase in colony forming units and up-regulation in the expression of mycobacterial resuscitation promoting factor genes. Present study thus suggests that some host or viral factors might be involved in the reactivation of dormant tubercle bacilli in spite of maintained CD4 counts which further needs to be deciphered extensively.

## Acknowledgement

We acknowledge Indian Council of Medical Research, New Delhi, India for partially funding the study.

## List of abbreviations

ADC: Albumin Dextrose Catalase
CFU: Colony Formimg Units
HIV: Human Immunodeficiency Virus
*M. luteus*: *Micrococcus luteus*
*M. tb*: *Mycobacterium tuberculosis*
Rpf: Resuscitation Promoting Factor
TB: Tuberculosis

